# ALaSCA: A novel *in silico* simulation platform to untangle biological pathway mechanisms, with a case study in Type 1 Diabetes progression

**DOI:** 10.1101/2023.03.16.532913

**Authors:** Carla Louw, Nina Truter, Wikus Bergh, Martine van den Heever, Shade Horn, Radouane Oudrhiri, Dawie van Niekerk, Ben Loos, Raminderpal Singh

## Abstract

**Introduction:** The analysis of signaling pathways is a cornerstone in clarifying the biological mechanisms involved in complex genetic disorders. These pathways have intricate topologies, and the existing methods that are used for the interpretation of these pathways, remain limited. We have therefore developed the Adaptable Large-Scale Causal Analysis (ALaSCA) computational platform, which uses causal analysis and counterfactual simulation techniques. ALaSCA offers the ability to simulate the outcome of a number of different hypotheses to gain insight into the complex dynamics of biological mechanisms prior to, or even without, wet lab experimentation. ALaSCA is offered as a proprietary Python library for bioinformaticians and data scientists to use in their life sciences workflows. Here we demonstrate the ability of ALaSCA to untangle the pivots and redundancies within biological pathways of various drivers of a specific phenotypic process. This is achieved by studying a major disease of global relevance, namely Type 1 Diabetes (T1D), and quantifying causal relationships between antioxidant proteins and T1D progression. ALaSCA is also benchmarked against standard associative analysis methods.

**Methods:** We use our *in silico* simulation platform, ALaSCA, to apply both a number of machine learning (ML) and data imputation techniques, and perform causal inference and counterfactual simulation. ALaSCA uses standard ML and causal analysis libraries as well as custom code developed for data imputation and counterfactual simulation. Counterfactual simulation is a method for simulating potential or hypothetical model outcomes in the field of causal analysis (Glymour, Pearl and Jewell, 2016). We apply ALaSCA to T1D by using proteomic data from Liu *et al.* (2018), as the patients were selected based on the presence of T1D susceptible HLA (human leukocyte antigen)-DR/DQ alleles through genotyping at birth and followed prospectively. The genetic cause of T1D in this cohort is therefore known and the mechanism and proteins through which it causes T1D are well-characterized. This biological mechanism was converted into a directed acyclic graph (DAG) for the subsequent causal analyses. The dataset was used to benchmark the causal inference and counterfactual simulation capabilities of ALaSCA.

**Results and discussion:** After data imputation of the Liu, *et al.* (2018) dataset, causal inference and counterfactual simulation were completed. The causal inference output of the HLA, antioxidant, and non-causal proteins showed that the HLA proteins had the overall strongest causal effects on T1D, with antioxidant proteins having the overall second largest causal effects on T1D. The non-causal proteins showed negligibly small effects on T1D in comparison with the HLA and antioxidant proteins. With counterfactual simulation we were able to replicate evidence for and gain understanding into the protective effect that antioxidant proteins, specifically Superoxide dismutase 1 (SOD1), have in T1D, a trend which is seen in literature. We were also able to replicate an unusual case from literature where antioxidant proteins, specifically Catalase, do not have a protective effect on T1D.

**Conclusion:** By analyzing the disease mechanism, with the inferred causal effects and counterfactual simulation, we identified the upstream HLA proteins, specifically the DR alpha chain and DR beta 4 chain proteins as causes of the protective effect of the antioxidant proteins on T1D. In contrast, through counterfactual simulation of the unusual case, in which the DR alpha chain and DR beta 4 chain proteins are not present in the model, we saw that the adverse effect which the antioxidant proteins have on T1D is due to the HLA protein, DQ beta 1 chain, and not the antioxidant proteins themselves. Future work would entail the application of the ALaSCA platform on various other diseases, and to integrate it into wet lab experimental design in a number of different biological study areas and topics.

## 1. Introduction

The analysis of signaling pathways is a cornerstone in clarifying the biological mechanisms involved in complex genetic disorders. Signaling pathways have intricate topologies, and exploratory wet-lab experimentation is generally used to elucidate the influence of different signaling components on disease outcomes. However, wet-lab experimentation is expensive in terms of laboratory space, cost of reagents, human resources and, moreover, time. In contrast, dry lab methods are capital-efficient and accessible. Due to the *in silico* nature of the work, where simulations are used to test hypotheses, dry lab methods surmount most of the limitations associated with wet lab experimentation. Additionally, *in silico* methods can greatly improve experimental design, by predicting potentially viable and non-viable outcomes before launching into costly and time-consuming experiments (Cárdenas, *et al.*, 2020). Therefore, effective and efficient *in silico* methods, to produce evidence around wet lab experimental design, are needed. This pertains to the field of metascience, a strongly emerging discipline concerned with increasing the quality and efficiency of scientific research (Metascience Symposium, Stanford University, 2019).

We have developed the Adaptable Large-Scale Causal Analysis (ALaSCA) software platform, which uses causal analysis and counterfactual simulation to untangle the pivots and redundancies within biological pathways. Pivots are key control points within a pathway which, when altered, can lead to significant change in the downstream processes. In contrast, redundancies in pathways are points that, when altered, have little to no effect on the phenotypic outcome of the pathway (Raimondi *et al.*, 2022., Stefanovski *et al.*, 2019).

ALaSCA consists of a pipeline for handling omics-type data, applying various machine learning and data imputation techniques, as well as causal inference and counterfactual simulation (Singh *et al.*, 2023). It has been developed based on custom code and also uses ML and causal analysis libraries which have been adopted by academia and industry as established standards for ML and causal analysis.

ALaSCA is based on formal Pearlian causal inference (PCI) techniques (Glymour, Pearl and Jewell, 2016., Pearl, 2000., Pearl and Mackenzie, 2018). Using PCI, we are able to quantify causal relationships using an array of different data types that span the hierarchy of biological organization from gene to phenotype. In comparison to other classic statistics techniques, ALaSCA generates actionable outcomes through simulations which are based on cause-and-effect relationships and not correlations from which causation is sometimes inferred. Other statistical or ML/AI methods often consider the relationships in biological mechanisms to be independent of each other, despite the complex nature of true biological mechanisms, and as such they are quantified independently when using these methods. In contrast, ALaSCA not only takes into account the causal relationships and their immediate surrounding network, the causal analyses and counterfactual simulations are performed in context of the entire biological mechanism.

Counterfactual, or ‘contrary-to-the-facts’ simulation, is a conviction based upon the human inclination to envision alternate outcomes to events that have already taken place (Glymour, Pearl and Jewell, 2016., Pearl, 2000). It can be used to simulate any model hypothesis, but is especially impactful in cases where the experimental procedure is difficult, not possible, or is unethical to conduct, or for retrospective simulation (Glymour, Pearl and Jewell, 2016., Höfler, 2005). It is also useful for simulating fat-tailed distributions and black swans - events which are highly unlikely to occur, but can have high impact (Cirillo and Taleb, 2020., Taleb, 2007). Counterfactual simulation allows for the simulation of multi-causal models as is often seen in diseases, mental disorders, and biological processes having multiple causes. The complex nature of these processes is taken into account when counterfactual simulations are conducted. Therefore all subsequent “downstream” effects of an intervention can be simulated accurately, irrespective of the complexity of the cascade of effects (Höfler, 2005). PCI and counterfactual simulation have become widely accepted approaches for conducting causal inference in a number of fields including biology, epidemiology, and medicine (Hernán and Robins, 2020., Höfler, 2005., Imbens and Rubin, 2015). In this study, we use the ALaSCA platform to quantify and simulate causal relationships between HLA proteins, and Type 1 Diabetes (T1D) progression, as mediated via antioxidant proteins.

T1D is a well-characterized and -understood disease, with availability of rich datasets proving ideal for demonstrating the utility of computational methods. The recent emergence, and continuous improvement of novel computational methods, combined with the subsequent integration of experimental data and computational techniques, has enabled an unprecedented level of detailed molecular understanding of the mechanisms involved in biological processes (Cárdenas, *et al.*, 2020). It is accepted that the main cause of onset and progression of T1D is rooted in genetic factors, coupled with subsequent autoimmune reactions. However, much is still unknown with regards to the specific mechanisms and pathways which directly mediate the effectors involved. T1D is a polygenic disease, and one of the principle determining genetic factors in its incidence is the inheritance of mutant major histocompatibility complex II (MHC-II) alleles. In humans, the MHC complex is referred to as the HLA complex, with polymorphisms of the HLA genes being directly linked to T1D susceptible genotypes (Morran *et al.*, 2015). Susceptible, as well as protective DR-DQ haplotypes are present in all human populations. Specific HLA DR/DQ alleles [e.g., DRB1*03-DQB1*0201 (DR3) or DRB1*04-DQB1*0302 (DR4), mediate the risk of T1DM progression, while HLA alleles such as DQB1*0602 are associated with dominant protection from T1DM in multiple populations (Erlich *et al.*, 2008). Our current understanding of the T1D mechanism originates from the failure of insulin producing beta- (β) cells maintaining tolerance to specific β-cell antigens (Figure 1; adapted from (KEGG, 2023., Erlich *et al.*, 2008)). When certain antigen-presenting cells (APC), such as pancreatic-resident dendritic cells and macrophages, erroneously process insulin-producing β-cells, it ultimately results in the destruction of these cells. Post-processing, β-cell proteins present on the surface of the APC, in a complex with MHC-II molecules, whereafter immunogenic signals from APC activate CD4+ T cells, mainly of the Th1 subgroup. These Th1 cells then produce IL-2 and IFNγ, which activate macrophages and cytotoxic CD8+ T cells. Activated macrophages and cytotoxic CD8+ T cells destroy β-cells through one or both of the following mechanisms: 1) direct interactions between antigen-specific cytotoxic T cells and a β-cell autoantigen-MHC-I complex on the β-cell, and/or 2) certain non-specific inflammatory mediators, in particular cytokines (IL-1, TNFα, TNFβ, IFN), and subsequently, IL-, and/or IFN- and/or TNF-induced free radicals/oxidants, resulting in oxidative stress. Three known cellular antioxidant defense systems operate in the mammalian system to reduce oxidative stress. This includes 1) low-molecular-mass antioxidants, 2) sequestration and repairing of systems and 3) antioxidant enzymes. Whilst the systems are interdependent, we focus on the system employing antioxidant enzymes to underwrite the trend shown in literature, of antioxidant proteins exerting a protective effect on T1D progression (Lei and Vatamaniuk, 2011).

**Figure 1:**
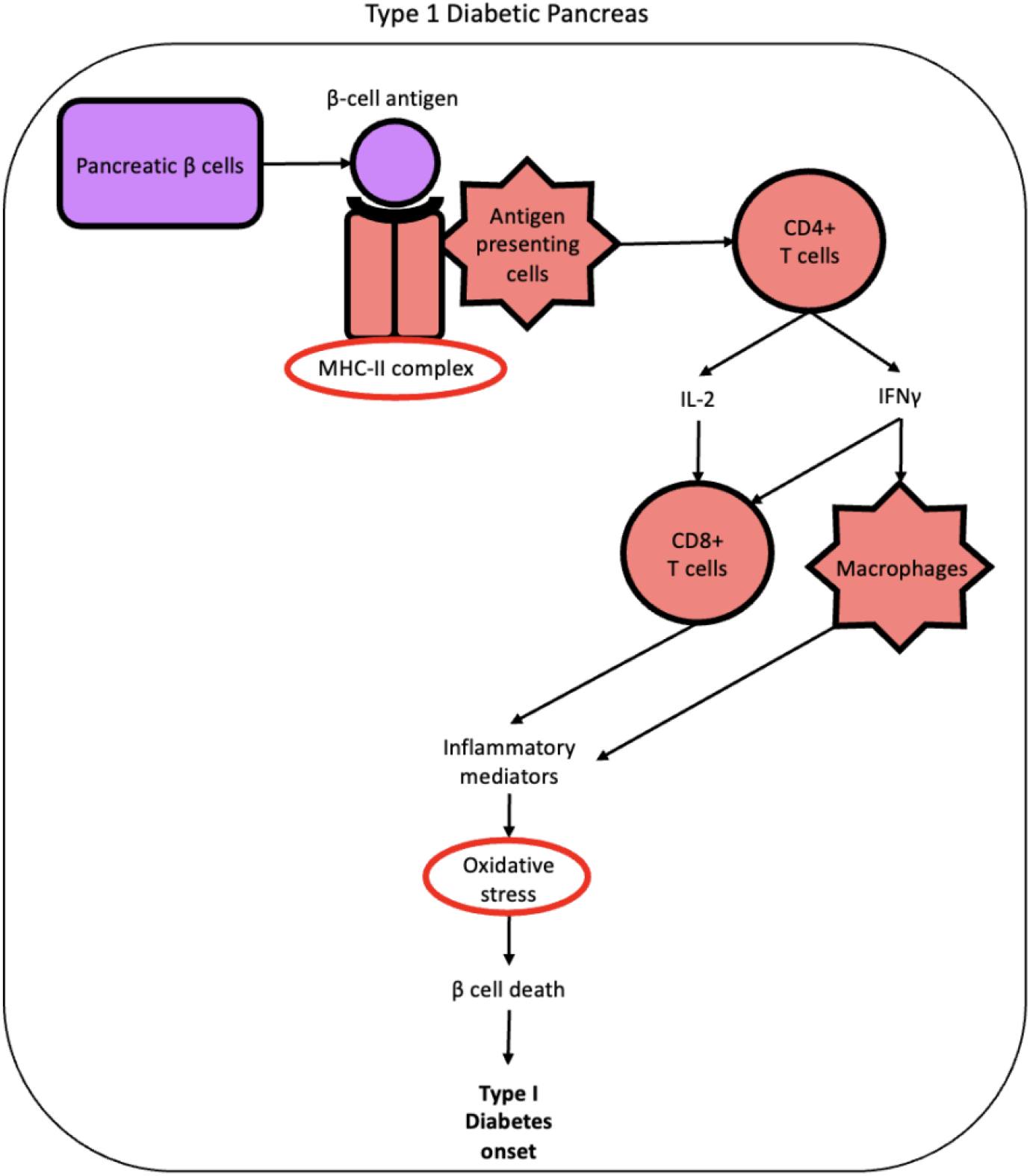
A well-accepted disease mechanism of T1D, including the MHC-II complex (circled in red) and resultant oxidative stress and T1D onset, adapted from KEGG (2023) and Erlich *et al.* (2008).

In the Methods section of this paper, we first provide an overview of the key components in the ALaSCA platform, as well as the T1D study and dataset used. We then, in the Results and Discussion section, describe step-by-step how the ALaSCA platform was applied to the T1D data and show significant findings from each step, focusing on the application of its causal analysis and counterfactual simulation capabilities. Finally, in the Conclusion section we highlight the insights into the T1D mechanism gained from the ALaSCA output.

Through the use of ALaSCA, we have untangled the dynamics within the T1D disease mechanism as represented in this study. We have identified which pathway branches mediate either an inhibitory or adverse (increasing) effect on T1D progression and how the HLA proteins affect T1D via the antioxidant proteins. This shows ALaSCA’s ability to untangle the pivots and redundancies within biological pathways of various drivers of a specific phenotypic process.

## 2. Methods

### 2.1 Key components in the ALaSCA platform

The ALaSCA *in silico* experimentation platform (Singh *et al.*, 2023) is a novel computational pipeline that consists of an array of algorithms for the processing and analysis of omics-type data followed by causal inference and counterfactual simulation. The counterfactual simulation functionality allows us to run multiple hypotheses for the simulation of various wet lab experiments *in silico*. ALaSCA supports the reduction of the cost of exploratory wet lab experimentation by aiding in the experimental design process and providing evidence for the true causal relationships in biological mechanisms.

The ALaSCA platform is a proprietary Python library, which consists of four major sections:

I. Data cleaning, formatting, and imputation
II. DAG creation from the biological mechanism, and de-risking
III. Causal inference
IV. Counterfactual simulation

These sections are discussed in detail below.

#### 2.1.1 Data cleaning, formatting and imputation

The data cleaning and formatting steps employ standard methods, as used in the fields of machine learning and statistics (Bishop and Nasrabadi, 2006., Hastie, Tibshirani and Friedman, 2009). ALaSCA is built using Python 3 and open-source python libraries, such as Numpy, Pandas, and Scipy, among others (Harris *et al.*, 2020., McKinney, 2010., Van Rossum and Drake, 2009., Virtanen *et al.*, 2020).

A number of standard exploratory data analysis (EDA) and machine learning algorithms form part of ALaSCA’s data cleaning and formatting step which precede the data imputation step. These include classic EDA for the handling of anomalies and outliers in the data, as well as the machine learning techniques of clustering, t-SNE plot analysis, and principal component analysis (Bishop and Nasrabadi, 2006., Pedregosa *et al.*, 2011). These techniques are used in combination with the classic EDA for thorough data exploration before any further imputation or causal analyses are conducted.

The data imputation process uses one of the classes of non-parametric Bayesian regression algorithms, Gaussian Process Regression (GPR), also known as Kriging. This algorithm uses prior distribution in the data to determine the predicted outcome with a distribution in uncertainty (Bishop and Nasrabadi, 2006., Xiong, 2022). We have developed a novel function which is built around the GPR function of the open-source python library, Scikit-learn (Pedregosa *et al.*, 2011). Our function is used for the hyperparameter optimization of the GPR function which minimizes the variance in omics data. This novel functionality returns an imputed omics dataset which mimics the natural variance seen in biological data (Singh *et al.*, 2023., Truter *et al.*, 2022).

#### 2.1.2 DAG creation from the biological mechanism, and de-risking

##### 2.1.2.1 Directed Acyclic Graph (DAG)

Directed acyclic graphs are the visual representation of the causal diagram used during causal inference and counterfactual simulation. A DAG consists of nodes, which are representative of the biological components in the mechanism, such as proteins, genes, or other features. The nodes in the DAG are connected by directed edges, which represent the relationships between the nodes. These directed edges are indicative of a causal effect and are quantified during the inference process.

##### 2.1.2.2 DAGs in the ALaSCA platform

ALaSCA does not consider the relationships in a system independently, but takes into account the causal relationships in the context of the network of effectors that constitute a biological pathway. This returns output which is relevant to the system and the simulations are representative of the true nature of the biological mechanism.

Our causal analysis pipeline starts with the gathering of prior knowledge, which is one of the building blocks of Bayesian analyses. Prior knowledge - e.g. a summary of biological mechanisms with assumptions and relevant study data - of the biological system or disease we wish to study, is gathered via an extensive literature search and discussions with key opinion leaders (KOLs) in said field of disease or system. The prior knowledge is represented in the DAG structure. If the mechanism is not well-known, we develop the most likely view (or views if we test multiple hypotheses) of the biological or disease mechanism based on our literature search and discussions with KOLs. This mechanism is then translated into a DAG.

##### 2.1.2.3 A method for de-risking the use of published literature and study data

Using prior knowledge and data in life sciences is often risky as there are significant biases and omissions:

1. Papers are positively biased to ensure that significant results are published. In some cases this can lead to the omission of “insignificant” data or to data dredging where researchers can report misleading analysis results to ensure significant findings, sometimes without assessing the validity of their methods (Joober *et al.*, 2012).
2. In line with positively biased publications, negative data is rarely published. This is mainly due to the probable rejection from journals, but also due to “prior” knowledge where a researcher will assume that their negative results are incorrect.
3. Life sciences coincide with uncertainty. Moreover, even topics that have a large amount of research attached to them, can have gaps in knowledge that might have been inferred to be unnecessary or redundant by previous researchers.
4. Omissions can affect presentation of data and the author’s inferences. An author might decide to omit data from a biological sample that did not correspond to trends that was seen in the data from the other biological samples. An author might also omit certain inferences, whether that be consciously or unknowingly. When a certain hypothesis is investigated with the data, it is very possible that only a part of the data is being analyzed or interpreted, precluding other possible inferences.
5. In many fields of research there is a lack of data generation standards as well as data sharing standards. Data generation should be replicable from published methods and inferences reproducible from published analyses, and this is not always the case (Diaba-Nuhoho and Ampnsah-Offeh, 2021). Furthermore, many papers have no raw data attached to them and the results can only be assumed to be correct.
6. Research is dependent on funding biases. Whether a proposal for funding will be accepted depends on if the proposal makes sense, but also if it is in line with the direction of research that the funding institution believes is correct (the dominant hypothesis). This belief normally stems from previous research in the field, even if that research has not proven to be fruitful (yet). An example of funding bias is the continuous and preferred funding towards projects focusing on the amyloid hypothesis (Begley, 2019).

In general, data in life sciences can be analysed with one of two approaches, namely associative or correlative analyses (which include classic ML and AI algorithms), or Bayesian approaches (which include causal inference).

Associative approaches create additional risk as the complex network of causation in real-world systems is not exposed. Some biases and risks in classical (associative) methods include (Cawley and Talbot, 2010., Baer and Kalmanath, 2017):

1. The sampled data does not capture the data generative process.
2. Confirmation bias in the data selected.
3. Overfitting of a complex model to the noise and random fluctuation in the data.
4. There is little insight into the underlying mechanism generating the data.
5. Only the patterns that are present in the data are considered and not any other, possibly important, factors not captured in the data.

Bayesian methods and causal inference approaches also include some residual risk. These approaches create risk because they require prior biological assumptions, and therefore require structured risk mitigation methods to avoid GIGO (garbage in, garbage out). There are four key biases and risks that need to be addressed in an effective causal analysis platform focused on biomedical data:

1. Incorrect assumptions about the prior biological knowledge
2. Limited scope and flexibility of the hypothesized mechanism
3. Confirmation bias in the literature review
4. Study meta-data information being incomplete

ALaSCA’s risk mitigation method addresses all four of these, by creating multiple DAGs with different levels of assumptions and by employing three iterative loops (data and experience-driven). Figure 2 describes ALaSCA’s process of DAG creation and de-risking, starting with the DAG construction from biological understanding, and biological and causal rules. The causal question and the dataset are used to constrain the DAG and quantify the causal relationships in the DAG. The DAG is de-risked with the use of three iterative loops.

**Figure 2:**
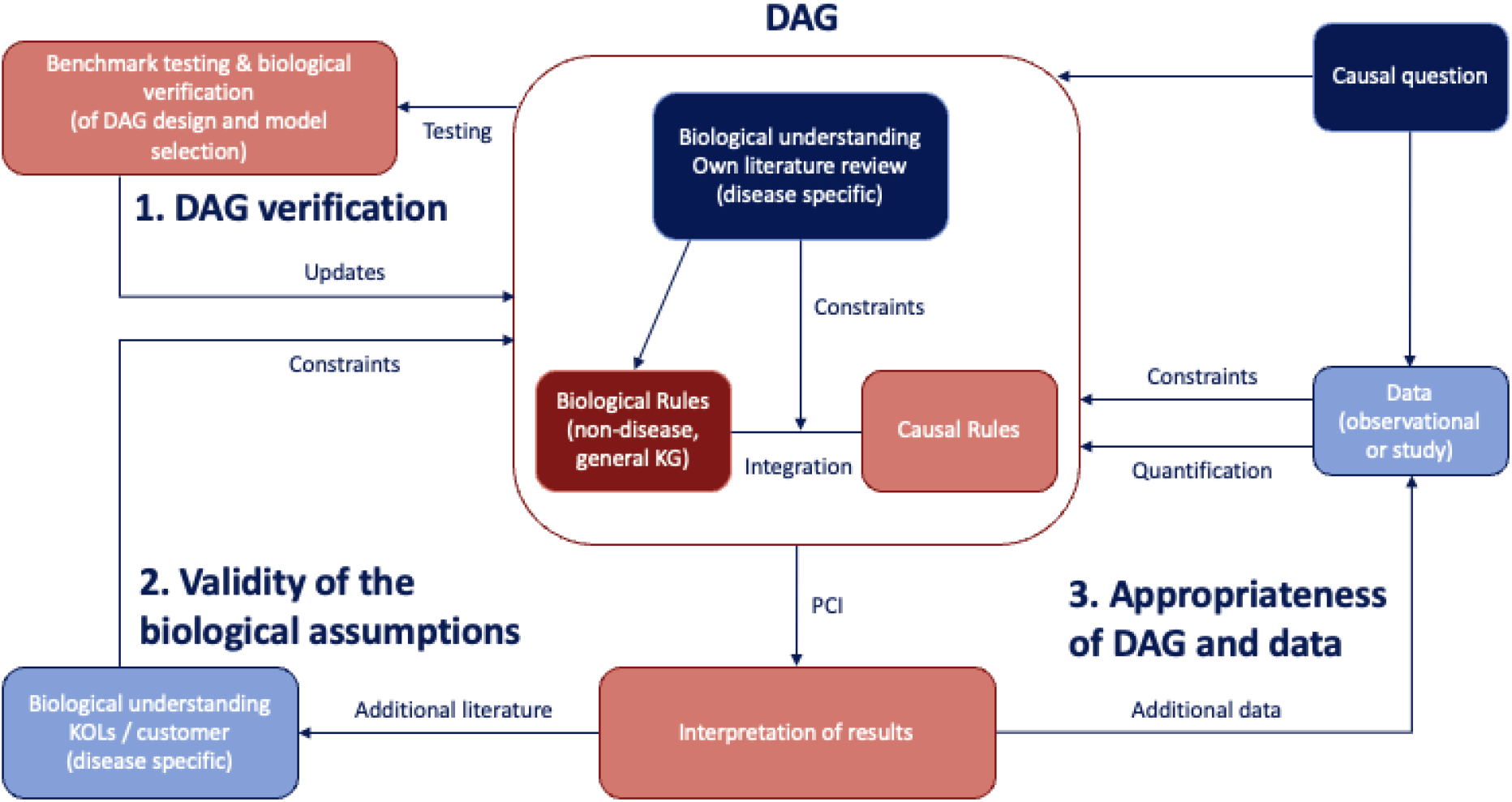
ALaSCA’s DAG creation and de-risking process. A DAG is constructed from biological understanding, and biological and causal rules. The de-risking process consists of three iterative loops: 1) DAG verification, 2) checking if the biological assumptions are valid, and 3) checking the relevance of the data for the DAG. The causal question as well as the data are used to constrain and quantify the causal effects in the DAG.

Below are listed the methods that ALaSCA uses to address the four biases and risks mentioned above, with reference to the iterative loops in Figure 2:

I. Incorrect assumptions about the prior biological knowledge Method: Identify the incorrect assumptions during iterations of loop 2 (Figure 2). Additional biological understanding from literature review and KOLs is then added. This is a pseudo-automated process leveraging scripts and peer-review processes. Example: Use study data to quantify the causal effects in a hypothesized DAG, if the study data is correct then any anomalies (i.e. unfeasible results) would most likely be due to the DAG structure being incorrect (i.e. incorrect assumptions).
II. Limited scope and flexibility of the hypothesized mechanism Method: Check for the appropriateness of fit of the DAG and the data during iterations of loop 3 in Figure 2, and highlight where the DAG requires improvement or which other datasets are required. Through iterations of loops 1 and 2 (Figure 2), the DAG scope can be adapted and extended as necessary. This process involves scripts and manual learning currently. Example: Upon interpretation of the PCI output it might be seen that all the causal effects in the DAG could not be quantified due to a limitation in the dataset. One can then find an additional dataset with which to quantify the DAG. Or it might be seen that the DAG does not explain the mechanism properly if any anomalies in the PCI results are seen and the dataset is sufficient - the DAG then needs to be improved through iterations of loops 1 and 2.
III. Confirmation bias in the literature review Method: Check the appropriateness of the fit of the DAG and the data during iterations of loop 3 (Figure 2), as well as test whether the biological assumptions are biased during iterations of loop 2 (Figure 2). This will highlight any confirmation bias that was introduced in the DAG design process. Further iteration of loop 2 (Figure 2) is then used to eliminate the confirmation bias. Example: If the study data is fair and complete, then anomalies in the PCI results are indicative of an unsuitable DAG. Loop 2 can then be used to identify sections of the DAG that contain bias.
IV. Study meta-data information is incomplete Method: Confirm that the DAG structure is correct, given the data and the causal rules, by using iterative loops 1 and 3 (Figure 2). This identifies whether the study meta-data (e.g. which confounding factors are included in the study) is incomplete, which is then remedied with further iteration of loop 3 and adding of more data with complete meta-data (Figure 2). This process is rule-based and is mostly automated based on the theory of PCI. Example: With the confirmatory and benchmark testing of loop 1 one can test if the DAG structure is complete. If it is not, then one can use loop 3 to add more data with the correct and complete meta-data such that we can accurately constrain the DAG and then quantify the causal effects accurately.

##### 2.1.2.4 DAG verification

To address DAG verification we use a validation process which consists of standard benchmark testing (Sharma and Kiciman, 2020) as well as biological verification. This validation process is a validation of our ability to design the appropriate DAG needed to determine causal inference, as well as the ability to choose the correct accompanying causal inference model (Louw *et al.*, 2023). One method that we have adopted in our DAG verification is to estimate how well the data fits the DAG. For example, we use a non-parametric approach as these are Bayesian networks. We also use an ensemble of DAGs and estimate treatment effects from that ensemble. This helps address some of the potential inaccuracies in the DAGs.

#### 2.1.3. Causal inference

Pearlian Causal Inference (PCI) is used for all causal analysis functionality in ALaSCA (Glymour, Pearl and Jewell, 2016., Pearl, 2000). We use the imputed dataset, which was originally selected based on relevance to the mechanism, to quantify the causal effects in the DAG. An array of different data types can be used for causal inference, such as clinical and other observational data, omics-type study data, as well as combinations thereof.

Standard causal inference Python libraries, such as PyWhy (DoWhy) are built into ALaSCA for the calculation of these causal values (Blöbaum *et al.*, 2022., Sharma and Kiciman, 2020). ALaSCA runs on a secure, public virtual machine instance, such as Google Cloud (2023).

#### 2.1.4. Counterfactual simulation

Similar to ALaSCA’s process of calculating causal inference values, classic PCI is also used for the counterfactual simulation functionality in ALaSCA (Glymour, Pearl and Jewell, 2016., Pearl, 2000). Counterfactual simulations allow one to simulate alternative or hypothetical scenarios which were not present in the data. ALaSCA uses structural causal modelling for the process of counterfactual simulation. Each node in the DAG (Figure 3) is represented by an equation (Equations 1 - 3) which consists of all parent nodes and the causal effects that these nodes have on the particular node, as well as an exogenous variable which is representative of any external, unmeasurable factors which might affect the node (Glymour, Pearl and Jewell, 2016).

**Figure 3:**
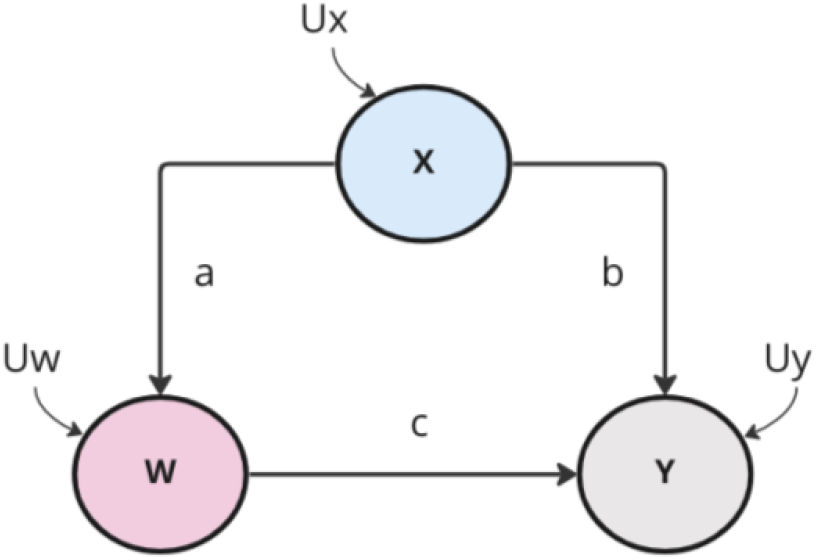
Example of a DAG with three nodes (W, X, and Y) and their respective causal effects (a, b, and c) as well as the exogenous variables of each node (U_W_, U_X_, and U_Y_)

Structural causal equations:

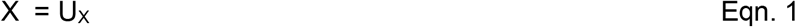

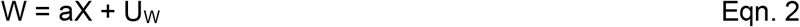

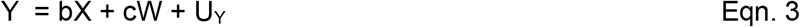

Here a, b, and c are the strength of the causal effects depicted as edges in the graph (Figure 3).

This process of counterfactual simulation consist of three steps as per classic PCI techniques:

I. Quantification of the exogenous variables (U_i_) and fixing of the exogenous variables to these values.
II. Modification of the model by intervening on a specific node (such as X) in the system and fixing the value thereof to a specific constant.
III. Predicting the effect of the counterfactual simulation on the outcome node (Y) by solving for the equation of Y given the updated values of U_i_ and X.

A custom Python library was developed for the counterfactual simulation in ALaSCA. This library allows us to run counterfactual simulation for a single intervention or multiple simultaneous interventions (Singh *et al.*, 2023).

### 2.2 Application of the ALaSCA platform on T1D mechanism

#### 2.2.1 Summary of public T1D study

The proteomes (2235 proteins) of 10 healthy controls and 11 T1D patients were measured across 9 time points, from birth to development of islet autoimmunity and overt T1D. The patients were selected based on the presence of T1D susceptible HLA (human leukocyte antigen)-DR/DQ alleles through genotyping at birth and followed prospectively. The genetic cause of T1D in this cohort is therefore known and the mechanism and proteins through which it causes T1D are well-characterized. In order to identify the molecular drivers of T1D, protein abundance data was used in our study, but gene expression (transcriptomic) data can also be used. The computational capability of calculating causal values can be used irrespective of whether the dataset is a transcriptomic dataset or the type used in our study (protein abundance).

Specific HLA proteins, as well as antioxidant genes and proteins, representative of the oxidative pathway, were included in the disease mechanism. Both of the selected genes and protein groups are known to contribute to T1D disease progression. A third protein group shown in Table 2 acts as a negative control; these proteins are associated with T1D, but are less likely to drive disease progression.

In this study the causal effects of proteins, known to be involved in the development of T1D (Table 1), on disease progression are determined using ALaSCA’s causal inference capability. Furthermore, these causal values are compared to those of proteins less likely to contribute to disease progression (Table 2). These proteins function as structural proteins and/or are involved in processes unrelated to the progression of T1D. The causal inference results of the Liu *et al.* (2018) study are used for subsequent counterfactual simulation of interventions on the T1D mechanism, to determine how the HLA proteins affect T1D progression via the antioxidant proteins.

**Table 1:**
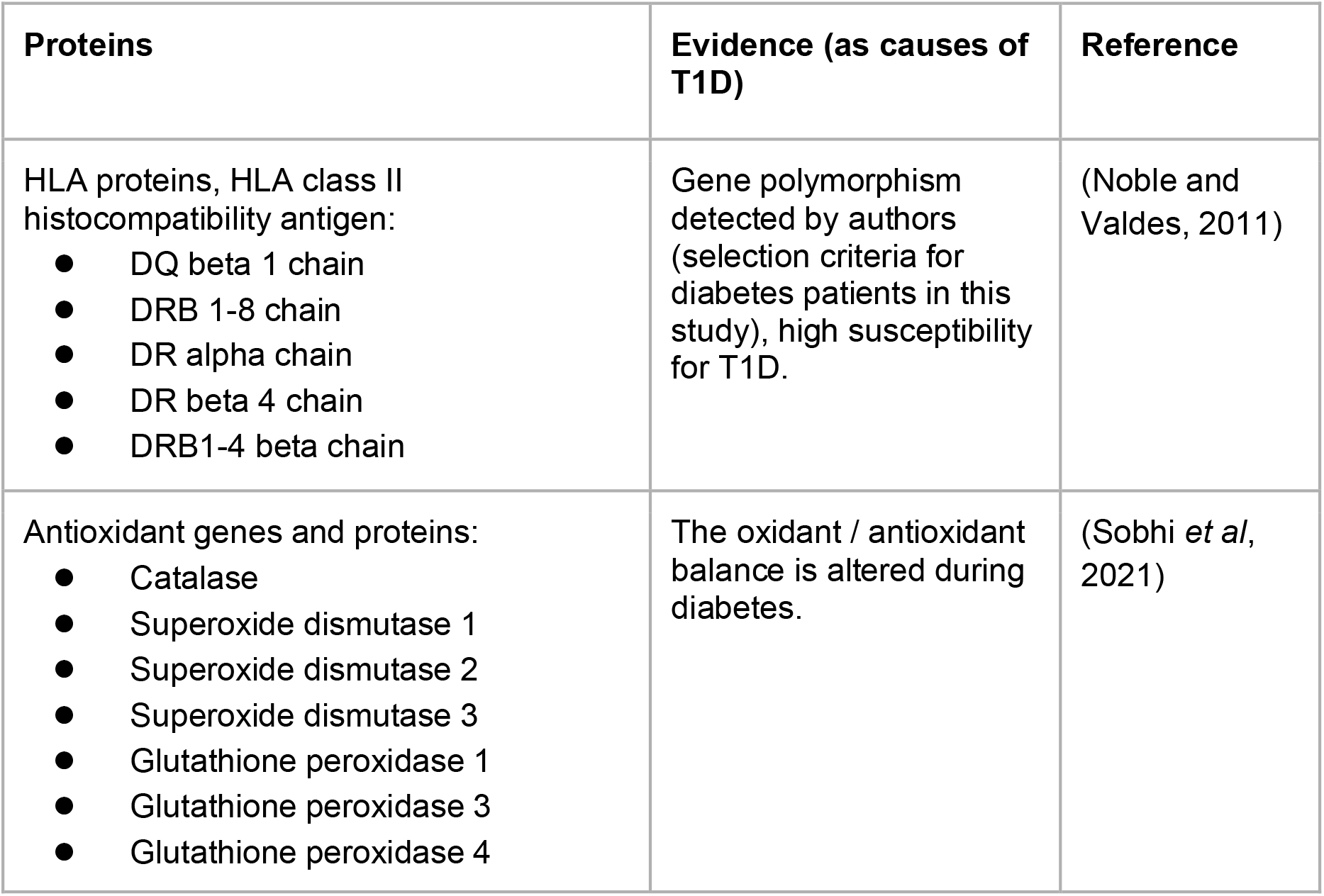
Proteins involved in driving T1D progression

**Table 2:**
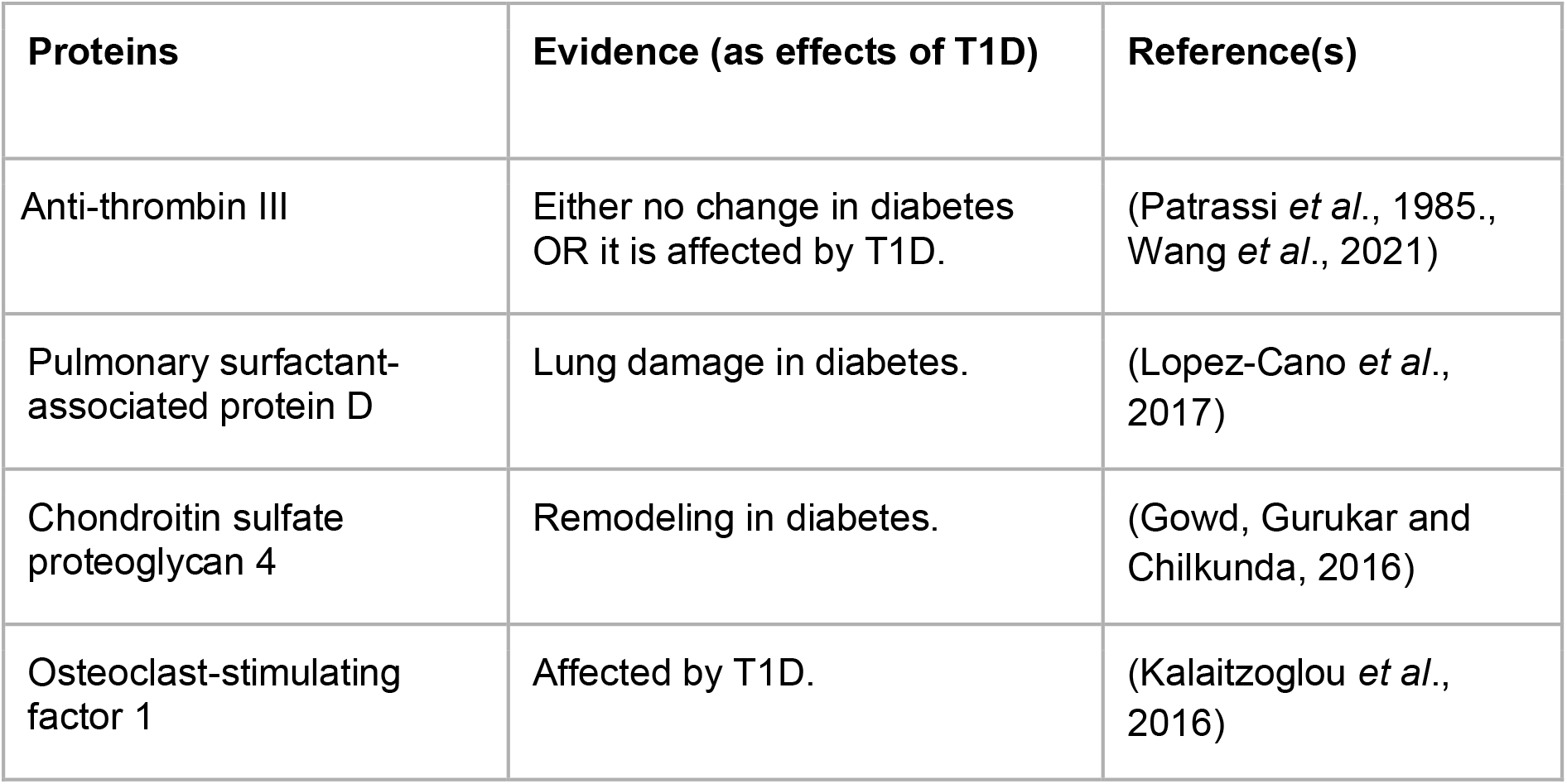
Proteins less likely to be engaged in T1D progression

| Proteins | Evidence (as effects of T1D) | Reference(s) |
| --- | --- | --- |
| Anti-thrombin III | Either no change in diabetes<br>OR it is affected by T1D. | (Patrassi *et al.*, 1985.,<br>Wang *et al.*, 2021) |
| Pulmonary surfactant-associated protein D | Lung damage in diabetes. | (Lopez-Cano *et al.*, 2017) |
| Chondroitin sulfate proteoglycan 4 | Remodeling in diabetes. | (Gowd, Gurukar and Chilkunda, 2016) |
| Osteoclast-stimulating factor 1 | Affected by T1D. | (Kalaitzoglou *et al.*, 2016) |

#### 2.2.2 Identifying true causal relationships instead of associative relationships

We show that associative and causal relationships of weak and strong protein drivers of T1D cannot be separated by ranking all 2235 proteins by the strength of their independent causal effect on T1D only (without context of the disease mechanism). We compare the output of simple, three-node DAGs (Figure 4(a)) with the output of the more complex biological mechanism (Figure 6). The simple, three-node DAGs only take the individual protein effects on disease into account, independent of all other proteins or the disease mechanism, such as the three-node DAG shown in Figure 4(b) for Superoxide dismutase 1 (SOD1).

**Figure 4:**
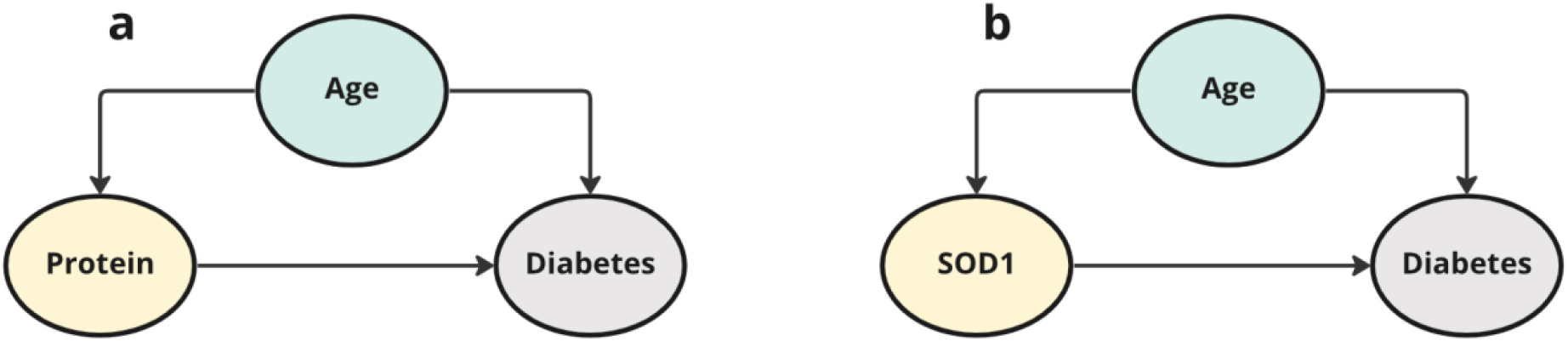
(a) The simple three-node DAG used to calculate the causal effect of each individual protein on T1D, independent of other proteins. (b) An example of this three-node DAG analysis applied to SOD1.

The use of three-node DAGs is expected to rank associative relationships between proteins and disease progression high, along with true causal relationships. As such, one would not be able to separate the proteins that are known to drive T1D (Table 1) from the proteins that are less likely to be involved in T1D but are affected by T1D (Table 2). The limitation of this approach (the use of simple three-node DAGs) is that it does not inherently include disease mechanism information in the constructed DAGs. Therefore proteins with high calculated causal effect values could either be drivers of T1D or proteins that are altered in T1D. The results are included in Supplementary Information A.

## 3. Results and discussion

### 3.1. Data cleaning, formatting and imputation of HLA, oxidative stress, and non-causal proteins

Data from Liu, *et al.* (2018) were cleaned and formatted, whereafter gaps in the data, between sampling events, were inferred using GPR. Figure 5 shows the imputed protein abundance levels of Superoxide dismutase 1 (SOD1) (Figure 5(a)) and Catalase (Figure 5(b)), both highlighted by Liu *et al.* (2018) as important in T1D for oxidative stress handling, for both control and diseased patients between the ages of 1 to 14 years. These imputed protein abundance levels of all proteins Tables 4(i) and 5(i) were used for all subsequent causal analyses and counterfactual simulations.

**Figure 5:**
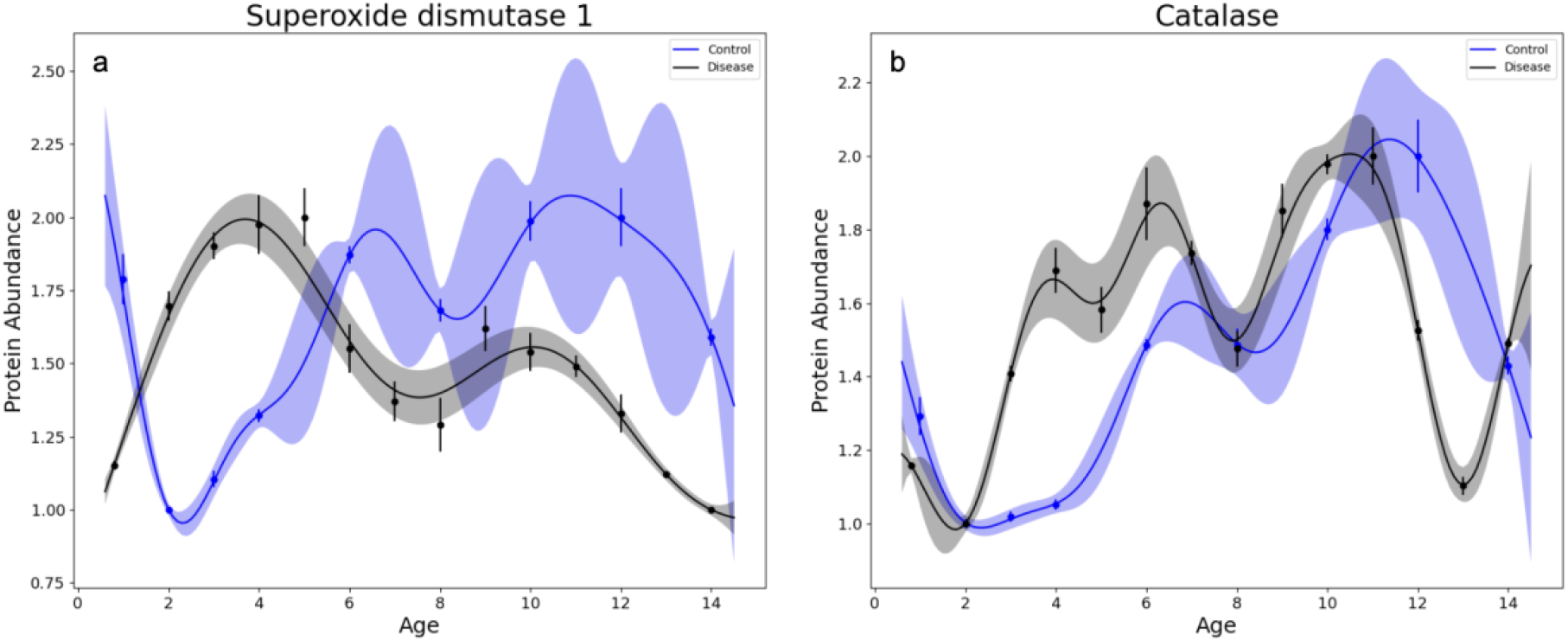
Inferred protein abundance using Gaussian Process Regression of the proteins (a) Superoxide dismutase 1 and (b) Catalase from the dataset of Liu, *et al.* (2018). The solid line and shaded area represent the mean and 95% credibility interval of the protein expression distribution, while markers with error bars show measurement means ± standard deviation. The blue and black lines represent the inferred levels of the protein in control and diseased patients respectively.

The GPR imputation does not alter the trends seen in the data, but merely imputes the gaps between the measured data points. This process assists in the subsequent causal inference analyses as well as interpretation of the original data.

From the GPR results, it is clear that the abundance of SOD1 increases early in the life of T1D patients (until age 4) (Figure 5(a)), whereas its abundance decreases as these patients age. This is in contrast with the control group, where SOD1 abundance increases from the age of 2 years. Catalase abundance seems to increase in both T1D patients and the control group over their lifespan (Figure 5(b)), until around age 11, however it increases more drastically in the T1D group between the ages of 2 and 6.

### 3.2. T1D disease mechanism to DAG

Given that the disease mechanism of T1D is well characterized, the process of translating the disease mechanism into a DAG was rooted in the knowledge from literature. The DAG developed for this study is also guided by the T1D study dataset used for quantifications thereof (Liu *et al.*, 2018). A DAG (Figure 6) is constructed from the disease mechanism. Alterations in Major Histocompatibility Complex Class II (MHC-II) are modelled through HLA proteins, while alterations in free radical metabolism are modelled through changes in abundance of antioxidant proteins involved in handling oxidative stress (Table 1). The DAG (Figure 6) is a visual representation of the pathways and protein-protein interactions through which these proteins are expected to drive T1D progression.

**Figure 6:**
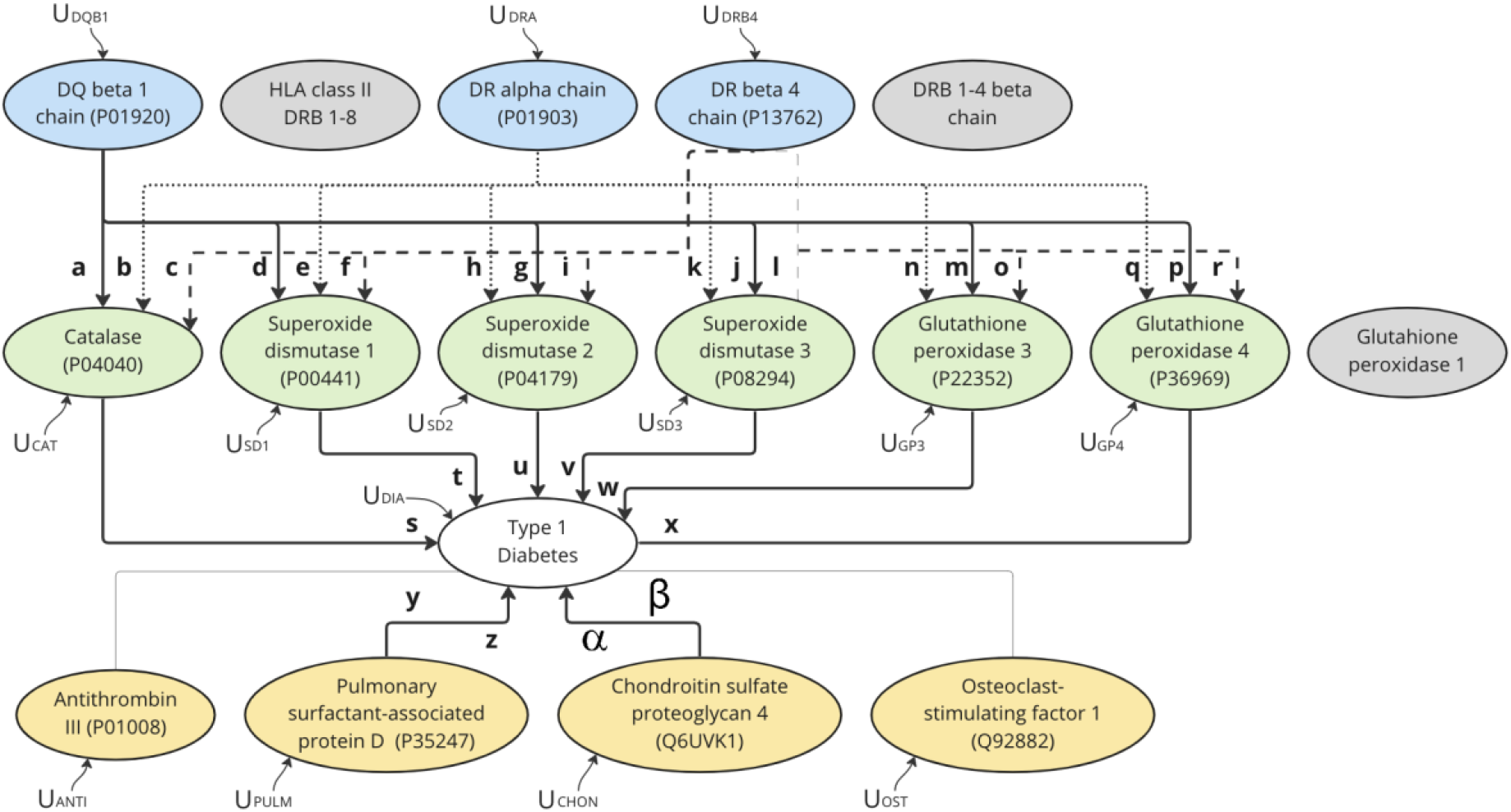
The mechanism DAG shows the HLA proteins (blue) which affect Type 1 Diabetes (T1D) via free radical metabolism, represented by the antioxidant proteins (green). Proteins for which either disease or control data could not be collected are shown in grey. Proteins less likely to be drivers of T1D are shown in yellow. The T1D disease progression is represented by the white node. The direct causal effects a - β are shown on the respective DAG edges, while the exogenous variable of each node is shown as Ui. Age (in years), not shown in the DAG, is a confounding factor for all proteins and T1D.

Figure 6 shows the DAG based on the mechanism of T1D, which contains all the proteins of Tables 1 and 2. The HLA and antioxidant proteins of Table 1 are shown in blue and green respectively with the HLA proteins affecting T1D via the antioxidant proteins. The grey nodes in the DAG indicate the proteins for which either disease or control data could not be collected. The direct causal effects between the proteins, or between the proteins and T1D, are represented by the symbols a - β and correspond with parameters in Equations 4 - 17 below. Similarly, the exogenous variables shown in Figure 6 (Ui) correspond with the exogenous variables in Equations 4 - 17.

### 3.3. Causal inference of HLA, oxidative stress, and non-causal proteins

The causal effect values in the disease mechanism DAG (Figure 6) were calculated with ALaSCA using the functions of the PyWhy (DoWhy) library (Blöbaum *et al.*, 2022., Sharma and Kiciman, 2020). The structural causal equations for this disease mechanism and the corresponding causal value estimates are shown as Equations 4 - 17 and Table 3 respectively.

**Table 3:** Estimated strength of effect values of the causal relationships, a - β, in the disease mechanism. The HLA, antioxidant proteins, and the non-causal proteins are shown in blue, green, and yellow, corresponding with the colors used in the DAG (Figure 6). The causal effects of the HLA proteins are generally higher than the other causal effects, while the causal effects of the non-causal proteins are, as expected, the smallest in comparison to the other two groups.

| HLA proteins on antioxidant proteins |  |  |  | Antioxidant proteins on T1D |  | Non-causal proteins on T1D |  |
| --- | --- | --- | --- | --- | --- | --- | --- |
| Causal effect | Strength of effect | Causal effect | Strength of effect | Causal effect | Strength of effect | Causal effect | Strength of effect |
| a | -30.86 | j | -0.16 | s | 0.012 | y | -0.0002 |
| b | -15.22 | k | 3.58 | t | -0.29 | z | -0.023 |
| c | -27.51 | l | 8.30 | u | -0.15 | $\alpha$ | 0.0037 |
| d | -3.08 | m | -8.93 | v | -0.071 | $\beta$ | -0.085 |
| e | -0.64 | n | -2.53 | w | -0.013 |  |  |
| f | -0.75 | o | 5.42 | x | -1.24 |  |  |
| g | -3.01 | p | -0.51 |  |  |  |  |
| h | 2.22 | q | 0.45 |  |  |  |  |
| i | 8.47 | r | 0.70 |  |  |  |  |

The parameters a - β in Equations 4 - 17 and Table 3 correspond to the parameters shown on the directed edges in Figure 6. These parameters are the strength of effect values of the causal relationships in the disease mechanism.

#### 3.3.1 Structural causal equations

The following abbreviations were used to represent the proteins in the structural causal equations: DQ beta 1 chain (DQB1), DR alpha chain (DRA), DR beta 4 chain (DRB4), Catalase (CAT), Superoxide dismutase 1 (SD1), Superoxide dismutase 2 (SD2), Superoxide dismutase 3 (SD3), Glutathione peroxidase 3 (GP3), Glutathione peroxidase 4 (GP4), Antithrombin (ANTI), Pulmonary surfactant-associated protein D (PULM), Chondroitin sulfate proteoglycan 4 (CHON), and Osteoclast-stimulating factor 1 (OST).

*HLA proteins:*

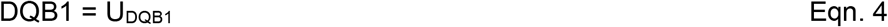

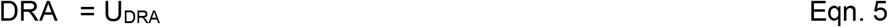

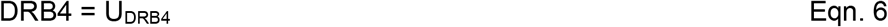

*Antioxidant proteins:*

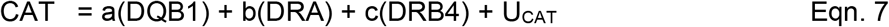

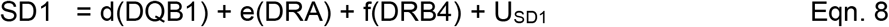

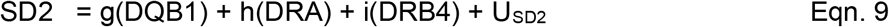

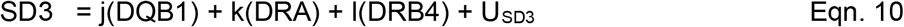

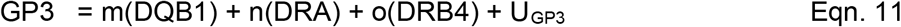

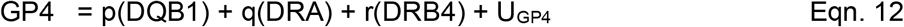

*Non-causal proteins:*

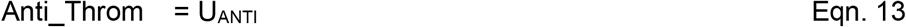

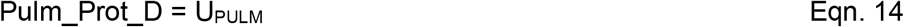

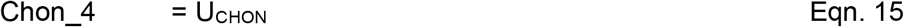

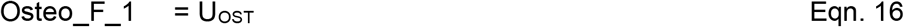

*Causal disease model outcome:*

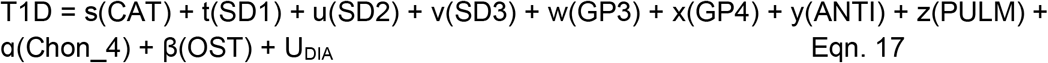

#### 3.3.2 Causal values

Table 3 shows that, on average, the proteins that are drivers of disease, both the HLA and antioxidant proteins (shown in blue and green respectively in Figure 6), have higher absolute causal values than the proteins less likely to be drivers of disease, the proteins shown in yellow in Figure 6. If a protein has a negative causal effect on another (or on T1D progression), it means that an increase in that protein will lead to a decrease in the other protein (or the disease outcome), and similarly a decrease will lead to an increase in the affected protein or disease outcome. Similarly, if a protein has a positive causal effect, an increase in the protein will have an increasing effect on the outcome, while a decrease in the protein will have a decreasing effect on the outcome.

When looking at both the change in protein abundance determined by GPR imputation (section 3.1) and the causal effect values of SOD1 and Catalase, a better understanding of their involvement in T1D progression is gained. The increases of SOD1 and Catalase observed in the T1D group in Figure 5 (section 3.1), is ascribed to the increase in oxidative stress before seroconversion (Liu *et al.*, 2018) and traditionally an increase in antioxidant abundance is regarded as a protective against oxidative damage in T1D (Lei and Vatamaniuk, 2011). This is supported by the negative causal value of SOD1 (t = −0.29) on T1D progression (Table 3). Unexpectedly, Catalase has a positive causal effect on T1D progression (s = 0.012; Table 3), which indicates that Catalase has an adverse effect on T1D. This anomaly is also observed in the study by Li, Chen and Epstein (2006) where an overexpression of Catalase in NOD mice resulted in the aggravation of T1D onset. To better understand the mechanisms behind SOD1 and Catalase’s effects on T1D, we performed counterfactual simulations using two studies in which their levels were altered (section 3.5).

### 3.4. Identifying true causal relationships instead of associative relationships

ALaSCAs ability to quantify causal relationships instead associative relationships was tested by comparing the causal output of the full disease mechanism (which is indicative of true causal relationships) with the causal output of simple, three-node DAGs (which are closer to associative analysis given the complex nature of biological mechanisms). We saw a difference between associative and causal relationships, as well as the importance of the effects of the surrounding mechanisms in the calculation of causal effects. See Supplementary Information A.

### 3.5. Counterfactual simulation of HLA, oxidative stress, and non-causal proteins

The three-step process of performing counterfactual simulation in classic PCI and the novel counterfactual simulation Python functionality of ALaSCA, which is based on this process, was used to simulate two specific scenarios. These two scenarios were taken from literature and were used as a validation of ALaSCA’s counterfactual simulation capability. The first scenario is the protective trend seen in antioxidant proteins as is summarized in the review by Lei and Vatamaniuk (2011), while the second scenario is the opposite, increasing effect that is seen in the study by Li, Chen and Epstein (2006). We also discuss why these two opposite trends are seen and how this is possible given that the disease mechanism is well understood.

#### 3.5.1 Simulating the protective effect of antioxidant proteins on progression of T1D

The structural causal equations (4 - 17) were used for the counterfactual simulation of a 50% increase in the antioxidant protein SOD1. The effect of this simulation on plasma antibody levels, which is used as a measurement of disease severity (i.e. in the place of the T1D node in Figure 6), is shown in Figure 7. The predicted antibody levels, with an increase in SOD1 are compared with the observed antibody levels as reported in the study by Liu, *et al.* (2018). The increase in abundance of this antioxidant protein has a decreasing effect on disease severity. This is the same qualitative trend in literature as is reported by Lei and Vatamaniuk (2011).

**Figure 7:**
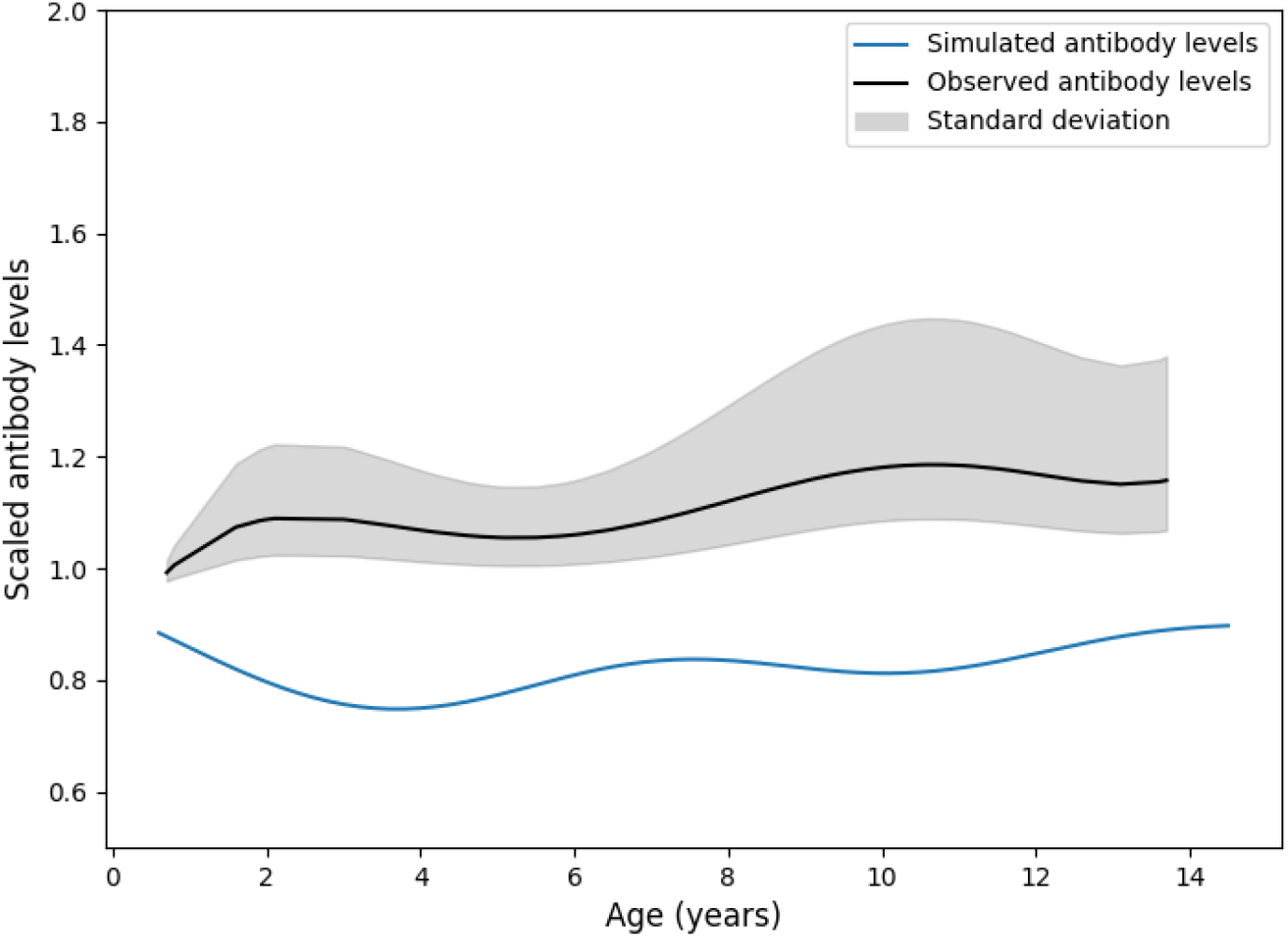
Comparison of antibody levels between the simulated T1D disease model with SOD1 activation (150% abundance) (blue) and observed antibody levels from literature (black) (Liu *et al.*, 2018). Simulation of the full disease mechanism model shows that the increase in abundance of Superoxide dismutase 1 protein has a decreasing effect on T1D (represented by antibody levels). This is similar to the trend seen in literature (Lei and Vatamaniuk, 2011).

#### 3.5.2 Simulating the adverse effect of antioxidant proteins on T1D

The review by Lei and Vatamaniuk (2011) highlights that the study by Li, Chen and Epstein (2006) is contradictory to the trend seen in literature with the latter showing that antioxidant proteins, specifically SOD1, have a positive effect on T1D, meaning that an increase in these proteins increase disease severity. To simulate the effects of this specific case, the structural causal equations (4 - 17) were adjusted to represent the model used by Li, Chen and Epstein (2006). This animal model shows that two of the HLA proteins (DR alpha chain and DR beta 4 chain) are not present and as such Equations 7 - 12 was adjusted as follows to Equations 7b - 12b:

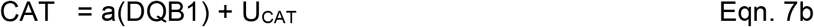

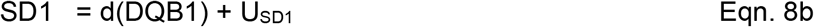

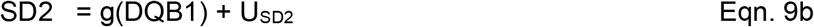

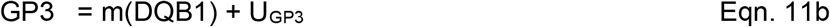

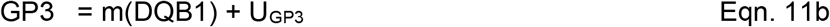

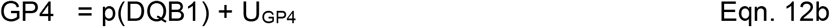

The remaining structural causal equations 4 - 6 and 13 - 17 were used with the updated equations 7b - 12b to simulate the counterfactual scenario with the effect of this simulation on scaled antibody levels shown in Figure 8. Li, Chen and Epstein (2006) reported that an increase in Catalase has an adverse effect on Diabetes. This same qualitative trend is seen in Figure 8 with an increase in this antioxidant protein leading to an increase in antibody levels.

**Figure 8:**
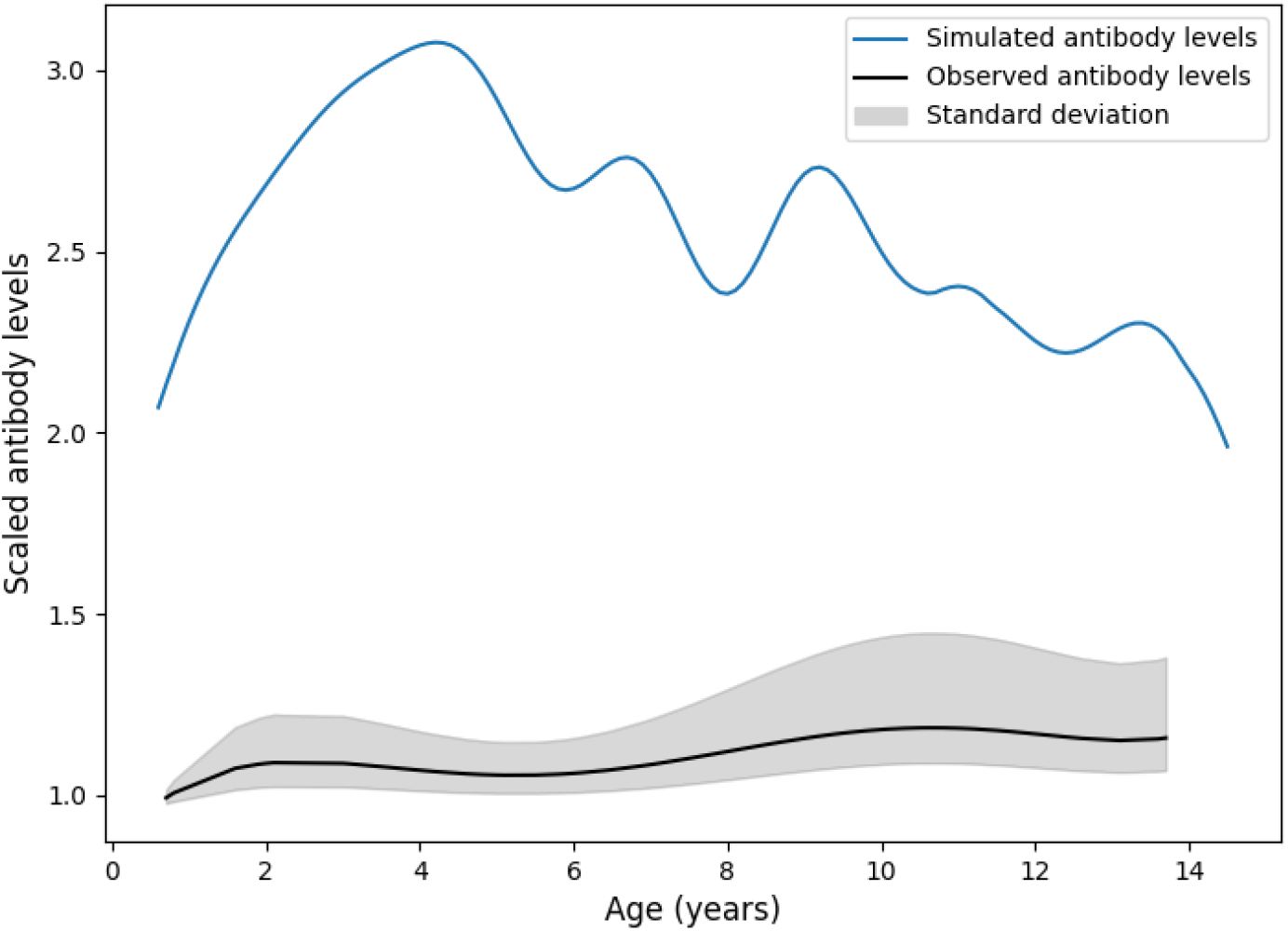
Comparison of antibody levels between the simulated T1D disease model with Catalase activation (150% abundance) (blue) and observed antibody levels from literature (black) (Liu *et al.*, 2018). Simulation of the Li, Chen and Epstein (2006) experimental model shows that the increase in abundance of Catalase has an increasing effect on Diabetes (antibody levels). The results are qualitatively comparable with the results seen in the Li, Chen and Epstein (2006) study.

### 3.6 Insight into the T1D mechanism

The reason that these two different trends are seen within the same disease mechanism can be explained by considering the causal effects. The main difference between the Li, Chen and Epstein (2006) study (Figure 8) and the rest of the field (Figure 7) is that the former study used the NOD T1D mouse model in which the DR alpha chain and DR beta 4 chain gene orthologs are not expressed and their associated proteins are therefore not present. The absence of these two HLA proteins leads to a significant change in the disease mechanism, as the HLA-DQ mouse ortholog contributes significant susceptibility to developing T1D in this model.

To further investigate these two above-mentioned trends, the total causal effects of the three HLA proteins present in the human disease mechanism (Figure 6) can be calculated with simple substitution using Equations 4 - 17. These total causal effects of the HLA proteins are shown in Supplementary Information B.

The total causal effect of DQ beta 1 chain on T1D has a positive value of 1.73423, while both the DR alpha chain and DR beta 4 chain proteins have negative total causal effects of −1.10933 and −2.91088, respectively. These total causal effects take into account all the pathways leading through the individual antioxidant proteins. The apparent protective effect that antioxidant proteins generally have on T1D (Lei and Vatamaniuk, 2011) is due to the effects that the upstream HLA proteins have on the antioxidant proteins and the subsequent effect that the antioxidant proteins have on T1D, which is an overall negative effect considering all three HLA proteins. Therefore, irrespective of which antioxidant protein is activated, an increase in antioxidant proteins will have a protective effect on T1D, given the larger disease mechanism (Figure 7).

In contrast, consider the case where the DR alpha chain and DR beta 4 chain proteins are not present in the disease mechanism, with DQ beta 1 chain being the only HLA protein present. The former two proteins are the drivers of the protective effect that is seen in the disease mechanism. Once these proteins are no longer present, only the positive effect of DQ beta 1 chain via the antioxidant proteins (in this case catalase), presides. Thus, an increase in catalase will now have the opposite effect to what is usually seen (Figure 8). This increase leads to an increase in disease severity, which corresponds to the findings of Li, Chen and Epstein (2006). However, from the analyses conducted on the disease mechanism we now see that this phenomenon is not due to the antioxidant proteins themselves, but the change in the disease mechanism as a whole, specifically the HLA proteins being the drivers of disease severity.

Both instances show that the effect of antioxidant proteins on T1D is largely driven by the upstream HLA proteins and cannot be ascribed to effects of the antioxidant proteins alone. This notion is confirmed by an observed chronological sequence of events of the major genetic determinants of T1D, originating on gene level - starting with polymorphisms of class II HLA genes, followed by their gene products, namely 1) HLA class I molecules, presenting endogenous antigens, and 2) HLA class II molecules presenting exogenous antigens to T cells, creating an HLA-peptide-T cell Receptor complex. This complex initiates the immune response, which in turn leads to oxidative stress and, thereafter, the production of antioxidant proteins to reduce oxidative stress (Noble and Valdes, 2011).

## 4. Conclusion

The capability of the ALaSCA platform to study the disease mechanism of a well-known disease, T1D, was tested. The known disease mechanism was adapted from KEGG (2023) and Erlich *et al*. (2008) into a DAG (Figure 6), which reflected the cause-and-effect type relationships in the mechanism, given the dataset used in this study (Liu *et al.*, 2018). The results of the causal inference and counterfactual simulation showed that the effect that antioxidant proteins have on T1D largely depends not on the antioxidant proteins themselves, but on the HLA proteins produced upstream due to polymorphisms of class II HLA genes. This was shown by using ALaSCA to replicate the trend seen in literature, where antioxidant proteins are shown to have a protective effect on T1D (Lei and Vatamaniuk, 2011) in Figure 7, as well as the unusual case where antioxidant proteins seem to increase T1D disease severity (Li, Chen and Epstein, 2006) in Figure 8. By analyzing the disease mechanism of both of these cases it is shown that the overall causal effect of the HLA proteins *via* the antioxidant proteins are the true causes of the protective or adverse effect seen in these studies, and not the direct effects of the antioxidant proteins on disease *per se.* In contrast, the study by Li, Chen and Epstein (2006) has an altered disease mechanism, where the only HLA protein present is the DQ beta 1 chain protein. In this altered disease mechanism the overall positive/increasing effect of DQ beta 1 chain is the only contributor to disease, although the presence of antioxidant proteins remain the same between both disease mechanisms.

In conclusion, we demonstrate ALaSCA’a ability to untangle the pivots and redundancies of the T1D biological pathway, showing that the effect of antioxidant proteins are dependent on the upstream HLA proteins and should not be considered to function independently thereof.

## 5. Future work

The ALaSCA platform is currently being applied to a variety of diseases and treatments in collaboration with our partners (biotechs, medtechs, etc.). In addition, we are actively working with large biopharma and traditional no-code platforms on integrating ALaSCA into their AI/bioinformatics workflows so that they can take advantage of causal analysis methods.

## 6. Acknowledgements

We would like to acknowledge Prof. Anthony Sedgwick (Founder of ThoughtDisruptor.com), Dr. Neil Wilkie (CEO of Mironid Ltd.), and Dr. Maksim Sipos (CTO of CausaLens Ltd.) for assistance with the review of this work.

## Notes

### Competing Interest Statement

Carla Louw, Nina Truter, Wikus Bergh, Martine van den Heever, Shade Horn, and Raminderpal Singh are employees of Incubate bio BV.

