## Supplementary Information A for "ALaSCA: A novel *in silico* simulation platform to untangle biological pathway mechanisms, with a case study in Type 1 Diabetes progression"

### Supplementary Information A: Identifying true causal relationships instead of associative relationships

ALaSCA's ability to differentiate between associative analysis and causal analysis was tested by comparing the causal output of the full disease mechanism (which is indicative of true causal relationships) with the causal output of simple, three-node DAGs (which are closer to associative analysis given the complex nature of biological mechanisms). Table 1(ii) shows the ranking of the proteins, known to be drivers of Type 1 Diabetes (T1D), out of 2235 proteins based on the causal values, while Table 2(ii) shows the ranking of the proteins less likely to drive T1D, but that may be affected by T1D. These causal values are calculated using simple, three-node DAGs (Figure 4) for each protein separately.

A high ranking of the proteins by their causal values is indicative of either a stronger causal effect (of the effect of the protein on disease progression) or a strong correlation between the protein and T1D, while a lower ranking is indicative of either a weaker causal value or weaker associative relationship. Healthy control or diabetic patient data were not collected for all proteins, for some proteins only disease or only control data were collected and as a result the causal effects of these proteins on disease progression could not be calculated. These proteins do not have a ranking in Tables 1(ii) and 2(ii), but the label "No data".

Although all of the HLA proteins are ranked higher than all other proteins in Tables 1(ii) and 2(ii), which is expected as these proteins are true causes of T1D, the antioxidant proteins, which also have causative effects on T1D, could not be separated from the proteins less likely to be involved in T1D (Table 2(ii)). The ranked causal values of Tables 1(ii) and 2(ii) show that a high correlative value is not necessarily indicative of a high causal value. As a result, nearly half (44%) of the proteins of Table 1(ii) could not be separated from (ranked higher than) Table 2(ii). For example, osteoclast-stimulating factor 1 (OSTF1) was ranked 541 out of 2235 proteins (in the top 25% of ranked proteins). OSTF1 is a protein that is affected by T1D due to enhancement of osteoclast activity, and not a known driver of T1D.

In contrast to the simple, three-node DAGs, Figure 6 shows the full disease mechanism DAG, which contains all the proteins of Tables 1(ii) and 2(ii). There are 18 (combined) causal effects that potentially drive T1D, calculated from the combined effects of the HLA proteins via the antioxidant proteins. These causal effects and their corresponding causal values are listed in Table 4. The proteins were ranked by their strength of causal effects. Of the 18 combined causal effects known to drive T1D (Table 4, section regarding Table 1(ii)), 15 causal effects ranked higher than the causal effects less likely to be involved in T1D (Table 4, section regarding Table 2(ii)).

ALaSCA's methodology of quantifying causal values in context of the entire biological or disease mechanism is able to distinguish between processes which are true drivers of disease and features which are less likely to be involved in disease, but might be associated with disease.

Table 1(ii): Ranking of proteins involved in driving T1D. All 2235 proteins in the dataset are ranked by their causal values. The proteins for which no data were included in the dataset are labelled as “No data”. Overall, the HLA proteins are ranked higher than the antioxidant proteins by their causal values.

| **Proteins** | **Evidence** | **Reference** | **Ranking** |
|---|---|---|---|
| HLA proteins, HLA class II histocompatibility antigen:<br>● DQ beta 1 chain<br>● DRB 1-8 chain<br>● DR alpha chain<br>● DR beta 4 chain<br>● DRB1-4 beta chain | Gene polymorphism detected by authors (selection criteria for diabetes patients in this study), high susceptibility for T1D. | (Noble and Valdes, 2011) | 161<br>No data<br>244<br>145<br>No data |
| Antioxidant genes and proteins:<br>● Catalase<br>● Superoxide dismutase 1<br>● Superoxide dismutase 2<br>● Superoxide dismutase 3<br>● Glutathione peroxidase 1<br>● Glutathione peroxidase 3<br>● Glutathione peroxidase 4 | The oxidant / antioxidant balance is altered during diabetes. | (Sobhi *et al*, 2021) | 1729<br>908<br>1222<br>1660<br>No data<br>1634<br>327 |

Table 2(ii): Ranking of proteins less likely to be engaged in T1D progression. All 2235 proteins in the dataset are ranked by their causal values. Overall, these proteins are ranked lower than the HLA proteins in Table 1(ii) by their causal values.

| **Proteins** | **Evidence** | **Reference(s)** | **Ranking** |
|---|---|---|---|
| Anti-thrombin III | Either no change in diabetes OR it is affected by T1D. | (Patrassi *et al*., 1985., Wang *et al*., 2021) | 1948 |
| Pulmonary surfactant-associated protein D | Lung damage in diabetes. | (Lopez-Cano *et al*., 2017) | 901 |
| Chondroitin sulfate proteoglycan 4 | Remodeling in diabetes. | (Gowd, Gurukar and Chilkunda, 2016) | 1619 |
| Osteoclast-stimulating factor 1 | Affected by T1D. | (Kalaitzoglou *et al*., 2016) | 541 |

Table 4: Ranking of the causal effects (by strength of causal effect) in the mechanism DAG by the causal values. Overall, the effects of HLA proteins on T1D, via the antioxidant proteins, are ranked higher than the effects of the non-causal proteins.

| | **Causal effect** | **Ranking** |
|---|---|---|
| HLA and antioxidant proteins known to drive T1D (Table 1(ii)) | DQ beta 1 chain -> Catalase -> T1D | 8 |
| | DQ beta 1 chain -> Superoxide dismutase 1 -> T1D | 2 |
| | DQ beta 1 chain -> Superoxide dismutase 2 -> T1D | 7 |
| | DQ beta 1 chain -> Superoxide dismutase 3 -> T1D | 20 |
| | DQ beta 1 chain -> Glutathione peroxidase 3 -> T1D | 15 |
| | DQ beta 1 chain -> Glutathione peroxidase 4 -> T1D | 4 |
| | DR alpha chain -> Catalase -> T1D | 14 |
| | DR alpha chain -> Superoxide dismutase 1 -> T1D | 13 |
| | DR alpha chain -> Superoxide dismutase 2 -> T1D | 9 |
| | DR alpha chain -> Superoxide dismutase 3 -> T1D | 11 |
| | DR alpha chain -> Glutathione peroxidase 3 -> T1D | 18 |
| | DR alpha chain -> Glutathione peroxidase 4 -> T1D | 6 |
| | DR beta 4 chain -> Catalase -> T1D | 10 |
| | DR beta 4 chain -> Superoxide dismutase 1 -> T1D | 12 |
| | DR beta 4 chain -> Superoxide dismutase 2 -> T1D | 1 |
| | DR beta 4 chain -> Superoxide dismutase 3 -> T1D | 5 |
| | DR beta 4 chain -> Glutathione peroxidase 3 -> T1D | 17 |
| | DR beta 4 chain -> Glutathione peroxidase 4 -> T1D | 3 |
| Proteins less likely to be engaged in T1D progression (Table 2(ii)) | Anti-thrombin III -> T1D | 22 |
| | Pulmonary surfactant-associated protein D -> T1D | 19 |
| | Chondroitin sulfate proteoglycan -> T1D | 21 |
| | Osteoclast-stimulating factor 1 -> T1D | 16 |
