## Supplementary Information B for "ALaSCA: A novel *in silico* simulation platform to untangle biological pathway mechanisms, with a case study in Type 1 Diabetes progression"

### Supplementary Information B: Total causal effects of HLA proteins

The total causal effects of the three HLA proteins present in the full disease mechanism (Figure 6) can be calculated with simple substitution using Equations 4 - 17:

*Total causal effect of DQB1 on T1D:*

$$
\begin{aligned}
T1D &= sa(DQB1) + td(DQB1) + ug(DQB1) + vj(DQB1) + wm(DQB1) + xp(DQB1) + U \\
&= (sa + td + ug + vj + wm + xp)(DQB1) + U \\
&= ((0.012)(-30.86) + (-0.29)(-3.08) + (-0.15)(-3.01) + (-0.071)(-0.16) + (-0.013)(-8.93) \\
&\qquad + (-1.24)(-0.51))(DQB1) + U \\
&= (-\ 0.37032 + 0.8932 + 0.4515 + 0.01136 + 0.11609 + 0.6324)(DQB1) + U \\
&= 1.73423(DQB1) + U
\end{aligned}
$$

Where the constant U is defined as $U = sb(U_{DRA}) + sc(U_{DRB4}) + sU_{CAT} + te(U_{DRA}) + tf(U_{DRB4}) + tU_{SD1} + uh(U_{DRA}) + ui(U_{DRB4}) + uU_{SD2} + vk(U_{DRA}) + vl(U_{DRB4}) + vU_{SD3} + wn(U_{DRA}) + wo(U_{DRB4}) + wU_{GP3} + xq(U_{DRA}) + xr(U_{DRB4}) + xU_{GP4} + y(U_{ANTI}) + z(U_{PULM}) + \alpha(U_{CHON}) + \beta(U_{OST}) + U_{DIA}$

*Total causal effect of DRA on T1D:*

$$
\begin{aligned}
T1D &= sb(DRA) + te(DRA) + uh(DRA) + vk(DRA) + wn(DRA) + xq(DRA) + U \\
&= (sb + te + uh + vk + wn + xq)(DRA) + U \\
&= ((0.012)(-15.22) + (-0.29)(-0.64) + (-0.15)(2.22) + (-0.071)(3.58) + (-0.013)(-2.53) + \\
&\qquad (-1.24)(0.45))(DRA) + U \\
&= (-\ 0.18264 + 0.1856 - 0.333 - 0.25418 + 0.03289 - 0.558)(DRA) + U \\
&= -\ 1.10933(DRA) + U
\end{aligned}
$$

Where the constant U is defined as $U = sa(U_{DQB1}) + sc(U_{DRB4}) + sU_{CAT} + td(U_{DQB1}) + tf(U_{DRB4}) + tU_{SD1} + ug(U_{DQB1}) + ui(U_{DRB4}) + uU_{SD2} + vj(U_{DQB1}) + vl(U_{DRB4}) + vU_{SD3} + wm(U_{DQB1}) + wo(U_{DRB4}) + wU_{GP3} + xp(U_{DQB1}) + xr(U_{DRB4}) + xU_{GP4} + y(U_{ANTI}) + z(U_{PULM}) + \alpha(U_{CHON}) + \beta(U_{OST}) + U_{DIA}$

*Total causal effect of DRB4 on T1D:*

$$
\begin{aligned}
T1D &= sc(DRB4) + tf(DRB4) + ui(DRB4) + vl(DRB4) + wo(DRB4) + xr(DRB4) + U \\
&= (sc + tf + ui + vl + wo + xr)(DRB4) + U \\
&= ((0.012)(-27.51) + (-0.29)(-0.75) + (-0.15)(8.47) + (-0.071)(8.30) + (-0.013)(5.42) + \\
&\qquad (-1.24)(0.70))(DRB4) + U \\
&= (-\ 0.33012 + 0.2175 - 1.2705 - 0.5893 - 0.07046 - 0.868)(DRB4) + U \\
&= -\ 2.91088(DRB4) + U
\end{aligned}
$$

Where the constant U is defined as $U = sa(U_{DQB1}) + sb(U_{DRA}) + sU_{CAT} + td(U_{DQB1}) + te(U_{DRA}) + tU_{SD1} + ug(U_{DQB1}) + uh(U_{DRA}) + uU_{SD2} + vj(U_{DQB1}) + vk(U_{DRA}) + vU_{SD3} + wm(U_{DQB1}) + wn(U_{DRA}) + wU_{GP3} + xp(U_{DQB1}) + xq(U_{DRA}) + xU_{GP4} + y(U_{ANTI}) + z(U_{PULM}) + \alpha(U_{CHON}) + \beta(U_{OST}) + U_{DIA}$
